# Incorporation of active cell-free expression lysates in chitosan coated alginate microcapsules

**DOI:** 10.64898/2026.06.04.730178

**Authors:** F. Merchant, F. Latifi, D. Sylaj, E. S. Wheeler, K. E. Loots, M. Coleman, V. Konjufca, S. Hoang-Phou

## Abstract

Oral routes of delivery are logistically simple and enables easy administration of therapeutics. However, oral delivery of proteins is still challenging due to the proteolytic environment within the gastrointestinal (GI) tract. To protect protein cargo from degradation, polymer encapsulation is commonly used, and when it is combined with cell-free gene expression (CFE) approaches that enable the rapid and flexible production of proteins, it potentially allows for on-demand production of protein therapeutics. Here, we investigated the suitability of chitosan coated alginate (Alg/Cht) microcapsules for encapsulation of proteins and CFE lysates for oral delivery. We show that CFE lysates can produce functional mCherry, a model fluorescent protein, in the presence of alginate polymers, although direct contact with chitosan did inhibit protein synthesis. We encapsulated CFE lysates or purified mCherry protein into alginate cores before crosslinking them using internal gelation techniques and coating with chitosan to test their protective capacity for oral delivery. Alg/Cht microcapsules protected mCherry protein cargo from degradation in simulated human gastric fluids and mouse gastric extracts and facilitated controlled cargo release upon exposure to conditions that simulate the intestinal environment. None of the individual CFE or encapsulation components induced inflammation in mouse GI tracts when administered via oral gavage. We also observed a delayed release of fluorescent bead cargo from Alg/Cht microcapsules in mouse intestines following oral gavage. Together, our data suggest that CFE lysate-loaded Alg/Cht formulations can be flexibly used to produce proteins and safely deliver them to the GI tract for potential therapeutic applications.

Graphical Abstract

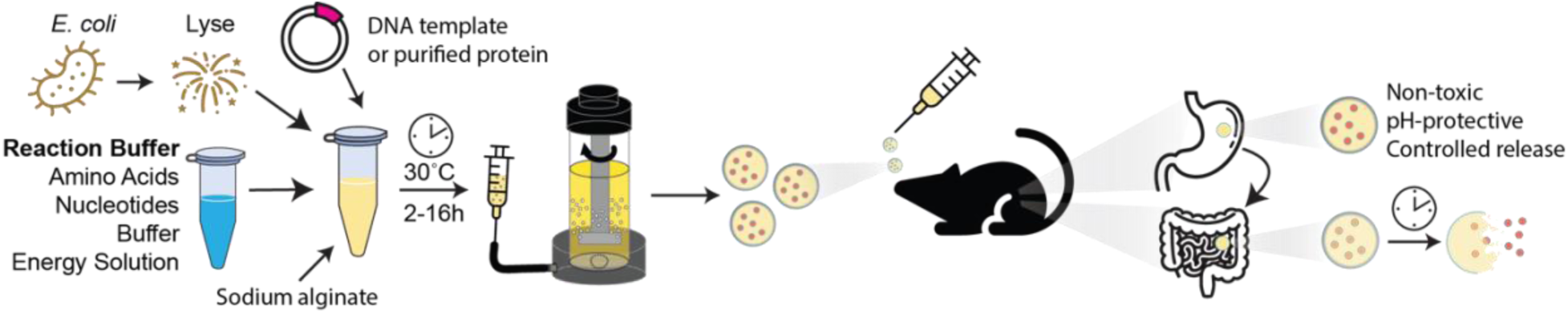

**Highlights:**

- Cell-free gene expression lysates are active in chitosan coated alginate (Alg/Cht) microcapsules.
- Alg/Cht microcapsules exhibit controlled release *in vitro* in simulated intestinal-like conditions.
- Cell-free and encapsulation components do not induce inflammation in the gastrointestinal tracts of male or female mice.
- Alg/Cht microcapsules show controlled delayed cargo release *in vivo* when orally gavaged in mice.

## Introduction

Oral therapeutics often have limited bioavailability due to the unique challenges associated with the varied environments of the gastrointestinal (GI) tract (Reviewed in [1]). Polymer encapsulation is a potential approach used to protect cargo for oral delivery and to promote controlled delayed cargo release and uptake in the GI tract. These formulations can take the form of a capsule or coating polymer, where an inner aqueous or semi-solid polymer “core” containing the cargo of interest is enclosed in a “shell” consisting of different polymers that often impart pH-protective or other stabilizing properties. Although there are many materials used for encapsulation of therapeutics and biologicals, alginate and chitosan are two natural polysaccharide polymers that predominate due to their limited toxicity and excellent biocompatibility [2].

Alginate is an anionic polymer consisting of linear chains of homo– or heteropolymeric blocks of α-L-guluronic and β-d-mannuronic acid, which can be crosslinked through their carboxyl groups through the introduction of divalent cations such as Ca^+2^ to form hydrogels (Reviewed in [3]). Chitosan is a linear cationic polymer derived from chitin where N-acetyl-D-glucosamine has been partially deacetylated to form β-1,4-D-glucosamine (Reviewed in [4]). Notably, alginate and chitosan spontaneously form stable structures through electrostatic interactions [5, 6], and chitosan coated alginate (Alg/Cht) particles show increased low-pH stability and controlled cargo release at neutral conditions. Thus, Alg/Cht formulations have demonstrated utility for oral delivery of small molecule drugs [7, 8] and other biological therapeutics, including hormones such as insulin [9] and whole cells for probiotic applications [10, 11]. However, encapsulation and oral delivery of recombinant proteins, which could be especially useful for immunization against pathogens, remains largely unexplored relative to other cargo materials.

Oral vaccines are easy to administer and can induce mucosal and systemic immunity which are critically important for protection against most pathogens. Live oral vaccines have been used for mass vaccination campaigns to curtail the spread of enteric diseases [12, 13]. However, live-attenuated pathogen formulations can suffer from drawbacks, exemplified by the oral polio vaccine (OPV), where OPV is crucial to quickly limiting outbreaks, but can also cause rare instances of vaccine associated paralytic polio, necessitating the careful consideration of strain formulations [14, 15]. Although orally available vaccines have significant potential due to their lower logistical burden, live vaccines still bear non-trivial risks and alternatives are needed. Recombinant protein-based subunit vaccines can bypass these risks as they are made in non-pathogenic prokaryotic or eukaryotic expression systems. However, protein antigens are especially vulnerable to the proteolytic conditions in the GI tract and their production and purification can be cumbersome.

Cell-free gene expression (CFE) is a technique that harnesses the protein production machinery extracted from cells, typically in a crude lysate, to enable the rapid production of proteins in a scalable and open system suitable for a variety of applications, including therapeutics and vaccine antigen production [16–18]. While CFE lysates can be made from both prokaryotic [19] and eukaryotic [20, 21] cells, *Escherichia coli*-based lysates are still the most widely used system (Reviewed in [22]), largely owing to their low barrier of entry and high protein yields. Endotoxin concentration is an important concern for bacterial or CFE produced recombinant protein vaccine antigens given systemically [23], and strategies to reduce endotoxin levels typically rely on stringent purification techniques or the use of low endotoxin *E. coli* strains [17, 24]. Oral delivery of *E. coli*-produced antigens is an attractive alternative that bypasses this constraint as the GI tract is naturally home to endotoxin-rich gut flora. However, potential degradation of orally administered protein antigens in the GI environment before reaching the gut-associated lymphoid tissues (GALT) may reduce their immunogenicity, necessitating the use of a variety of approaches to enhance antigen protection and uptake in the GALT. Importantly though, CFE lysates can be repackaged into novel form factors while maintaining their function, including encapsulated and hydrogel formats to modify their stability, lifetime, and durability [25–27].

Here, we sought to harness the ability of alginate core and chitosan shell encapsulation, generally referred hereafter as “Alg/Cht,” to enable oral delivery of proteins. We showed that CFE is active in the presence of polymer encapsulation materials in batch and encapsulated formats. Polymer encapsulation gave decreased overall protein production but still yielded enough protein typically used for vaccine dosages. Notably, none of the CFE or encapsulation formulation components induced inflammation or toxicity when given to mice through oral gavage. Lastly, we demonstrated that Alg/Cht encapsulation protected its protein cargo from low pH and controlled its delayed release at higher pH in simulated GI fluids and mouse GI fluid extracts, as well as enabled the release of fluorescent nanoparticles *in vivo* in mouse GI tracts, illustrating its potential for oral protein delivery applications.

## Results

### Cell-free gene expression lysates retained activity in the presence of polymers

Although cell-free gene expression (CFE) is a flexible and open system (Fig. 1A), lysate productivity can be sensitive to additives and other reagents, and we sought to determine whether they retained their activity in the presence of polymer encapsulation materials. To assay for CFE activity, we added the polymers or their solvents individually to cell free reactions making mCherry at a range of concentrations and measured the protein yield (Fig. 1B) and fluorescence output (Fig. 1C). We observed less mCherry protein yields by WB and reduced fluorescence in the reaction tubes with increasing concentrations of each material. Notably, we saw a near complete loss of mCherry production even at the lowest concentration (0.05%) of chitosan we tested, precluding its use as a core material, however it could still be used as a coating or “shell”, where it would have minimal contact with the CFE lysates [28]. As efficiency of CFE lysates is reduced in suboptimal conditions, we used the polymer concentrations at which there was still sufficient CFE protein synthesis activity to develop the encapsulation formulation, focusing on alginate concentrations between 0.5-1%.

**Figure 1.**
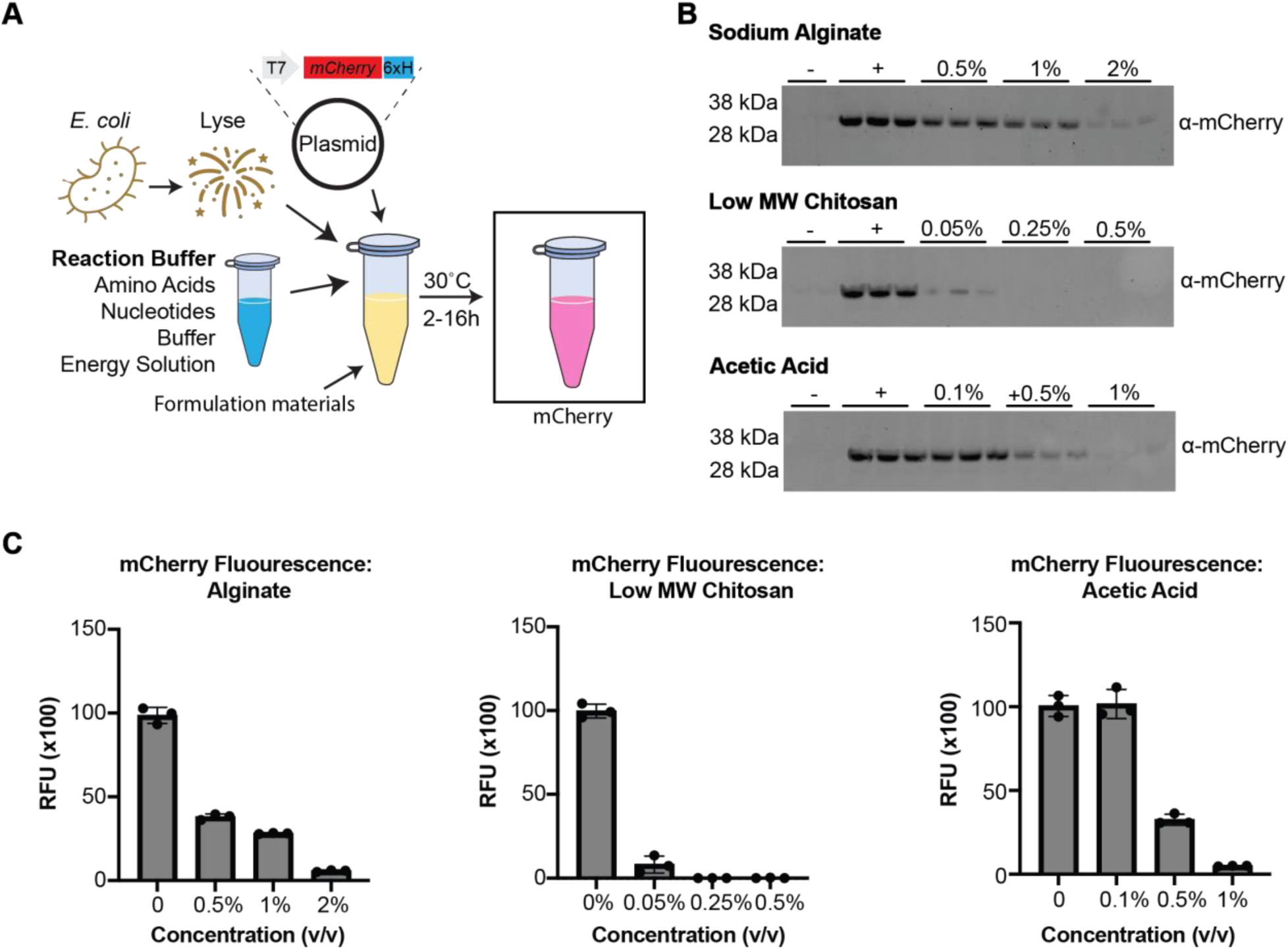
Cell-free lysates retain some protein synthesis activity in the presence of polymer materials. (**A**) Schematic of cell-free gene expression (CFE) of mCherry. (**B**) Western blots of CFE reactions containing increasing concentrations of polymer encapsulation materials probed with α-mCherry antibodies (**C)** Bar graphs depicting CFE mCherry fluorescence in the presence of sodium alginate (left), low MW chitosan (middle), and acetic acid (right).

### Cell-free lysates retained their activity after encapsulation in alginate

Since CFE lysates showed variable activity in the presence of alginate polymers in tube batch format, we wanted to confirm that activity is retained when lysates are encapsulated in droplet form. We performed initial alginate droplet production at macro-scale (millimeters) using manual syringe shearing into a single-step Ca^+2^-based external ionic gelation and chitosan coating bath to form macrocapsules (Fig. 2A). Encapsulation of CFE lysates with mCherry plasmid DNA in 0.5%-1.0% alginate cores with a 0.5% chitosan coat enabled the production of active mCherry protein after incubation overnight at 30°C in all alginate concentrations (Fig 2B). However, we observed low capsule structural integrity in the 0.5% alginate condition, where they were the most malleable and prone to leakage of the mCherry protein into the supernatant solution, thus we performed further tests using the 0.75% and 1.0% alginate formulations. To quantify the amount of mCherry produced in the capsule core (Fig. 2C), we produced Alg/Cht macrocapsules containing known concentrations of purified mCherry protein for a standard curve and compared its fluorescence to the mCherry produced in CFE lysate Alg/Cht macrocapsules. CFE lysate macrocapsules produced 0.14 +/− 0.01 mg/mL mCherry protein in the 0.75% alginate cores compared to 0.08 +/− 0.01 mg/mL in the 1.0% alginate cores (Fig. 2D). Although the 0.75% alginate formulation showed more leakage compared to 1.0% alginate, we reasoned that the increase in protein yield would be more than sufficient to overcome this difference. Therefore, all subsequent core formulations used 0.75% sodium alginate as a compromise between protein production yield and rigidity of the droplets.

**Figure 2.**
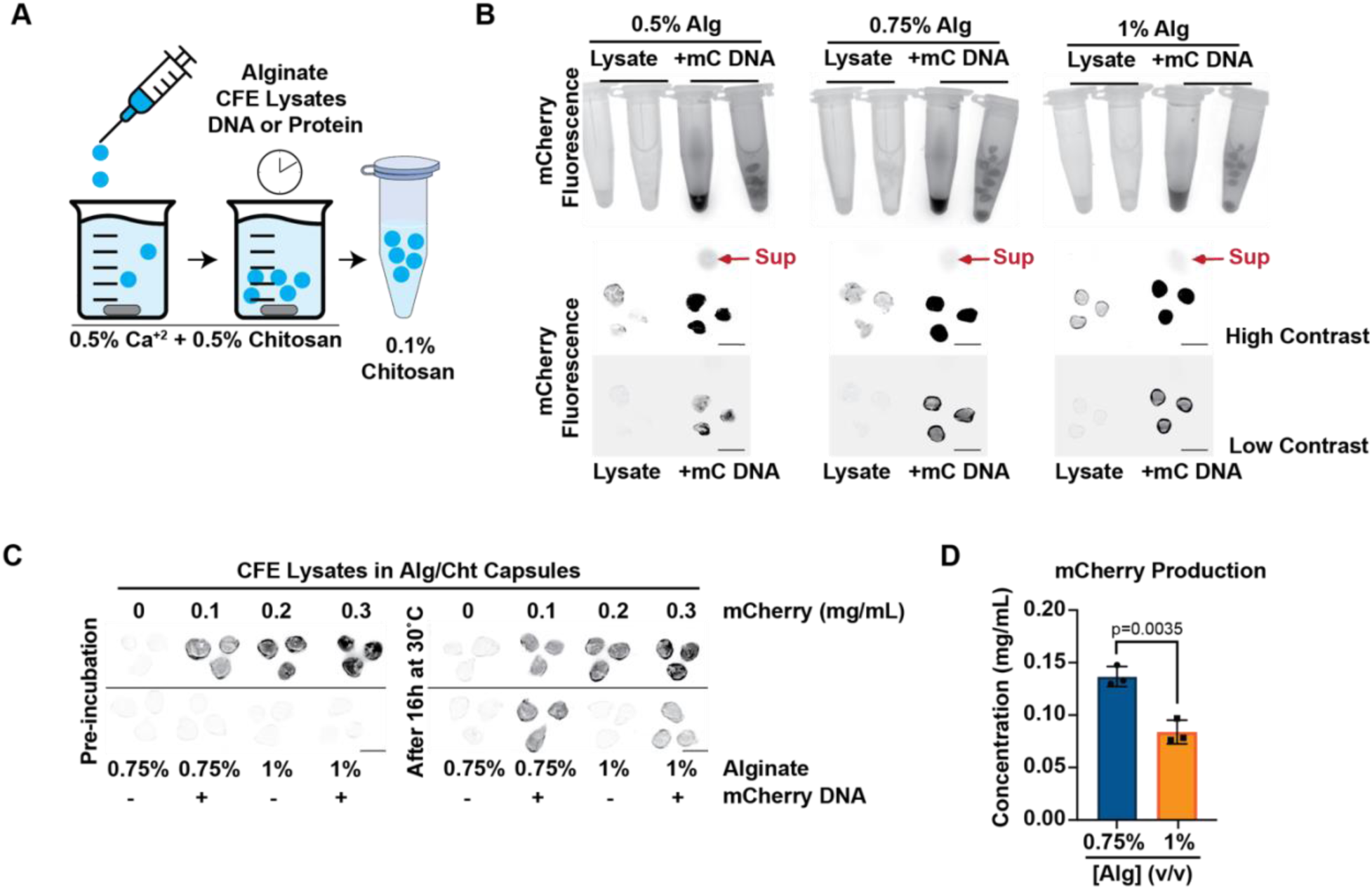
Cell-free gene expression activity is retained inside alginate macrocapsules after external ionic gelation. (**A**) Schematic of chitosan-coated sodium alginate (Alg/Cht) droplet generation. (**B**) Fluorescence scans of Alg/Cht capsules containing CFE mCherry in tubes (top) and on glass slides (bottom) at 520nm emission. Scale bar=5mm. Sup=Droplet supernatant. (**C**) Fluorescence scans of Alg/Cht capsules containing purified or CFE mCherry on glass slides at 520nm emission. Scale bar=5mm. (**D**) Bar chart showing yields of CF produced mCherry in Alg/Cht capsules using 0.75% and 1% alginate. n=3 independent experiments. *p*-value obtained using Student’s t-test.

### Cell-free protein synthesis activity was retained in water-in-oil emulsions but limited by acid-based internal ionic gelation

Although the macrocapsules created using manual syringe shearing maintained CFE lysate activity, they were too big for oral gavage administration to mice, which could impact their use as oral delivery vehicles. To decrease droplet size, we made a couple key changes to the droplet generation process. We switched to using the Micropore LDC-1 dispersion cell, which shears off droplets into a water-in-oil (W/O) emulsion after extrusion through 20µm pores (Fig. 3A). While this method generated droplets that met our requirements, we were unable to reliably generate stable microcapsules using the external ionic gelation method (not shown). However, we could recover stable microcapsules by switching to CaCO_3_-based internal gelation, which worked by incorporation of sequestered Ca^+2^ in the form of particulate CaCO_3_ in the core fluid and then release of that Ca^+2^ through acidification of the oil phase using 3.5% acetic acid (v/v).

**Figure 3.**
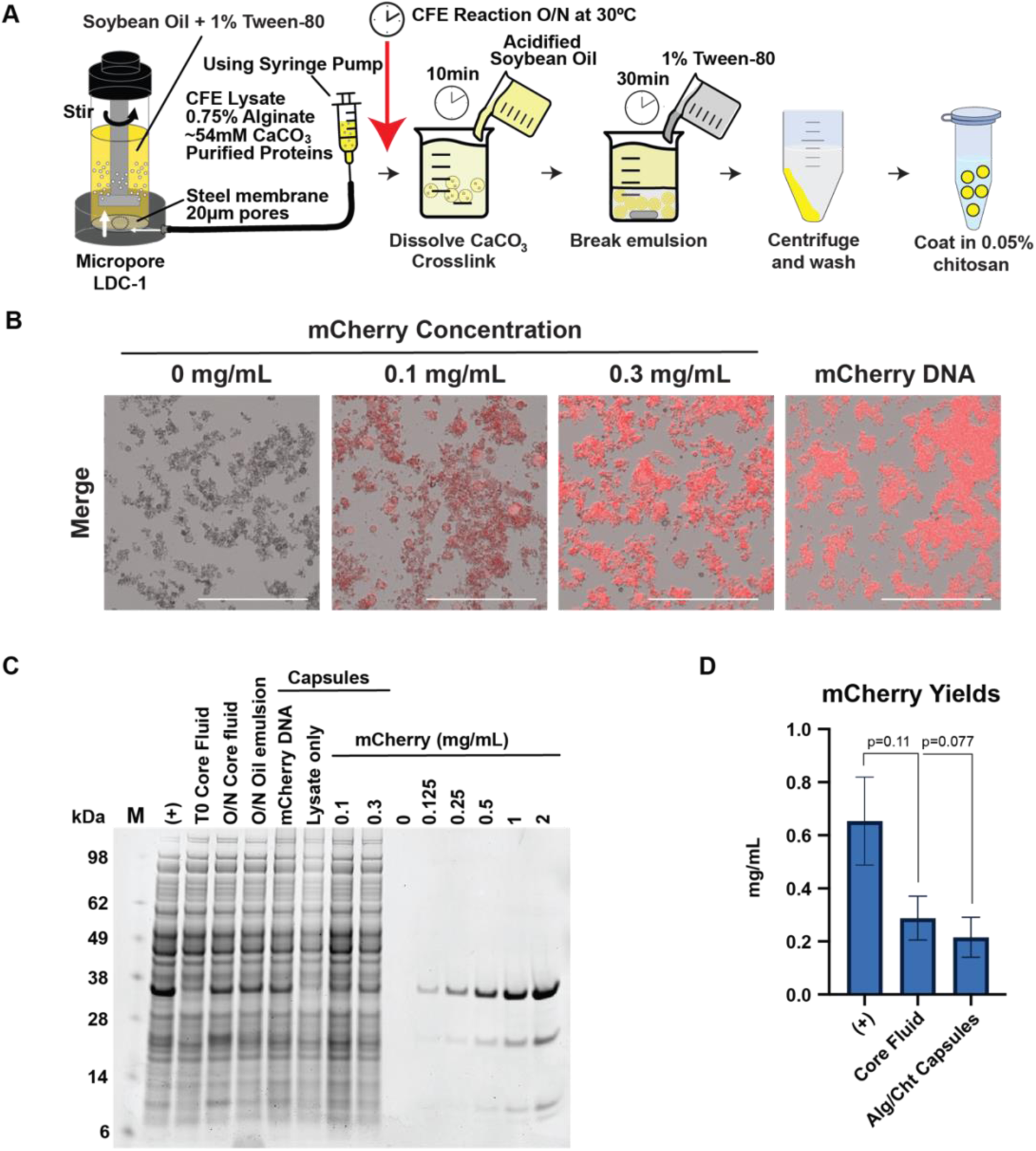
Cross-linking using acid-enabled calcium release limits cell-free gene expression in internally gelated alginate droplets. (**A**) Schematic of Alg/Cht droplet generation using the Micropore LDC-1 unit and internal gelation. Red arrow refers to droplet generation arrest and incubation to perform CFE reactions. (**B**) Fluorescence micrographs of Alg/Cht droplets containing purified mCherry protein or CFE mCherry. Scale bar=275µm. (**C**) Coomassie stained SDS-PAGE gel of CFE mCherry sampled across the droplet generation process. (**D**) Bar charts of CFE mCherry yields without additives (+), with alginate and CaCO_3_ (core fluid), or after microcapsule generation (coated droplets). n=2 independent experiments. *p*-values obtained using Student’s t-test.

We produced microcapsules using core fluids containing either CFE lysates + mCherry DNA or purified mCherry protein. Unfortunately, we observed a loss of CF-produced mCherry fluorescence after making Alg/Cht microcapsules using this method. Sampling the core fluid or droplets from CFE lysates at each step of the droplet generation process revealed a striking loss of mCherry fluorescence after internal gelation crosslinking of the alginate droplets (Supp. Fig. 1A). Interestingly, while mCherry fluorescence recovers in the mCherry protein-loaded sample after chitosan coating, it remains absent in the CFE lysate + DNA sample, suggesting that CFE activity is inhibited. Indeed, our preliminary results indicated that acetic acid strongly reduced the yield of mCherry protein (Fig. 1B) at the concentrations required to fully solubilize the CaCO_3_ used in the core fluid (Supp. Fig. 1B), corroborating the hypothesis that the acidification step inhibits the protein synthesis reaction.

To address this apparent incompatibility, we arrested the droplet production process after generating the W/O emulsion and incubated it overnight while shaking at 30°C to produce mCherry before continuing with crosslinking (Fig. 3A; red arrow). We observed active CFE-produced mCherry using this approach, with similar fluorescence intensities to the microcapsules loaded with 0.3mg/mL mCherry protein (Fig. 3B). SDS-PAGE analysis of the fluids or droplets from the production process against a purified mCherry standard curve revealed an upper-limit yield of 0.22 +/− 0.08 mg/mL mCherry protein in the final Alg/Cht microcapsules (Fig. 3C-D), which is roughly a 66% drop in yield compared to CFE lysates without any additives. Importantly, addition of 0.75% alginate to CFE lysates alone accounted for 56% of the drop in yield, in line with our earlier screening results, and suggested that processing the core fluid into W/O emulsions only modestly affected lysate productivity.

### Alginate/chitosan droplets exhibited controlled release in in vitro assays

An effective oral delivery vehicle for protein-based therapeutics must protect its cargo from the harsh proteolytic conditions of the stomach, resist large pH changes, and release its cargo at the desired location in the gastrointestinal (GI) tract. To test whether our formulation could fulfill these requirements, we produced and incubated mCherry Alg/Cht microcapsules in simulated human gastric and intestinal fluid (hSGF and hSIF, respectively; Table 1) [29] while shaking at 37°C in an *in vitro* release assay and took aliquots over time (Fig. 4A). To more closely mimic *in vivo* conditions at the stomach-proximal intestine junction, we mixed the droplet solution 1:1 (v/v) with hSIF after initial incubation with hSGF. Consistent with the idea that our Alg/Cht formulation can protect its mCherry cargo from denaturation and degradation in the stomach before releasing it in the more pH neutral intestine, we observed a sharp decline in mCherry fluorescence in the pelleted droplets (Supplementary Fig. S2A) with an inversely proportional increase in the supernatant only after introduction of hSIF (Fig. 4B-C). Western blotting of hSGF and hSIF microcapsule pellets and supernatants against the hexahistidine tag on mCherry using α-His antibodies confirmed the loss of mCherry protein from the capsules and increase in the supernatant after introduction of hSIF (Fig. 4D).

**Figure 4.**
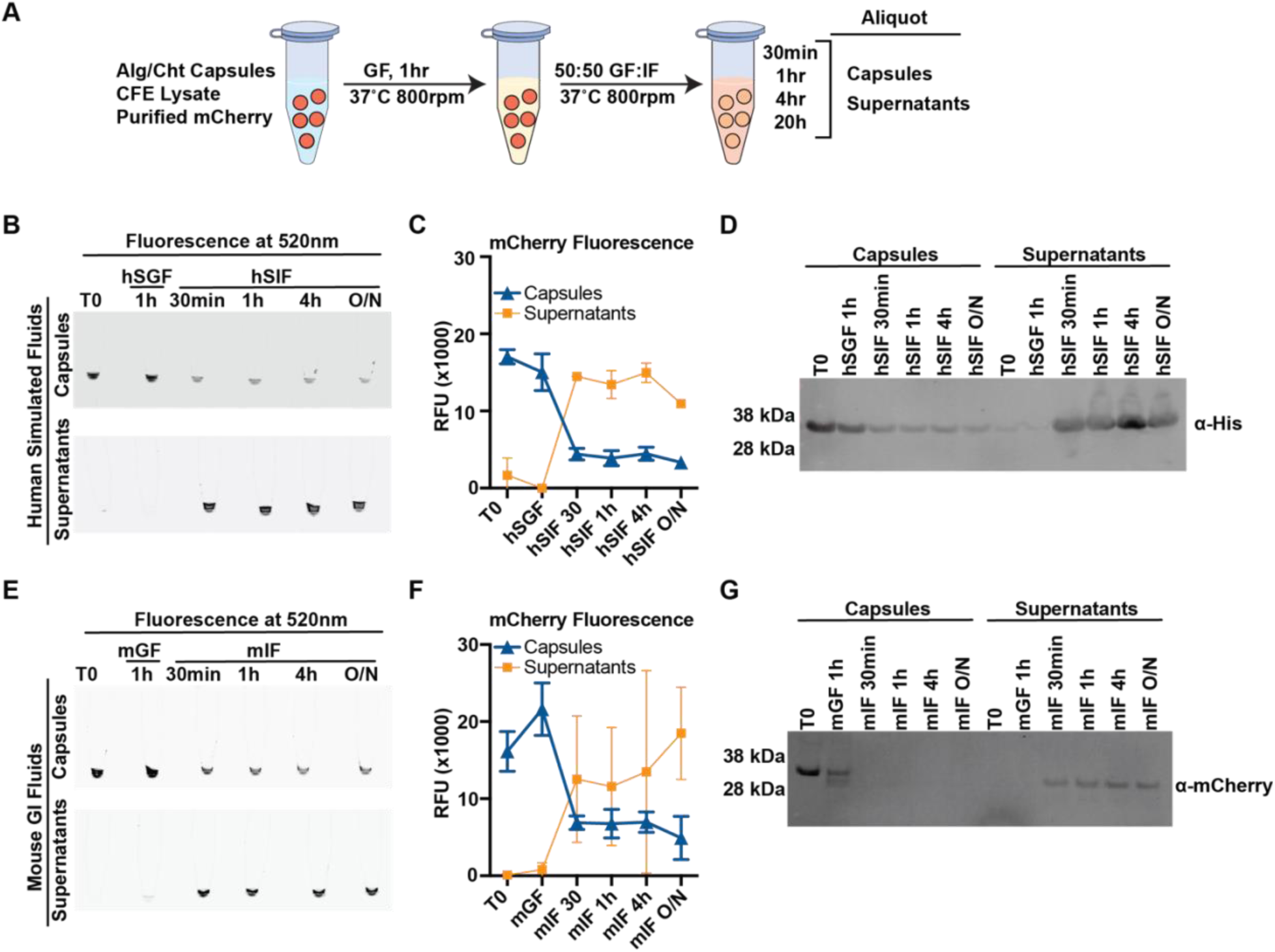
Chitosan coated alginate droplets allow controlled release of encapsulated protein in simulated intestinal conditions. (**A**) Schematic of experimental workflow for *in vitro* release testing. (**B**) Fluorescence scans of tubes containing mCherry Alg/Cht microcapsules and supernatants exposed to human simulated gastric or intestinal fluids (hSGF and hSIF, respectively) imaged at 520nm emission. (C) Line graph depicting quantification of fluorescence from (B). n=2 independent experiments. (D) Western blot of droplets or supernatants from (B) probed with α-His antibodies. (E) Fluorescence scans of tubes containing mCherry Alg/Cht microcapsules and supernatants exposed to mouse gastric or intestinal fluid extracts (mGF and mIF, respectively) imaged at 520nm emission. n=3 independent experiments. (F) Lline graph depicting quantification of fluorescence from (E). (G) Western blot of droplets or supernatants from (E) probed with α-mCherry antibodies.

While the results from human simulated fluids were promising, we wondered whether *bona fide* gastric and intestinal fluids would give similar results as they are more complex solutions. To validate our findings in a more physiological *in vitro* system, we extracted mouse gastric and intestinal fluids (mGF and mIF, respectively) and performed the same release assay. Pleasantly, we saw the same trends in mCherry release using mGF and mIF as we did for the hSGF and hSIF (Fig. 4E-F and supplementary Fig. S2B).

Interestingly, we could not detect mCherry through western blot with the same α-His antibodies used previously (not shown) after introduction of mIF. It is likely that the hexahistidine tag was getting cleaved from the mCherry protein through a thrombin cleavage site located between each sequence, and re-probing the membrane with α-mCherry antibodies enabled detection of the protein, showing similar trends to the simulated fluids (Fig. 4G).

### Oral delivery components did not induce inflammation in the GI tract

As we intended to use Alg/Cht microcapsules and bacterial lysates for oral delivery of protein-based therapeutics, we first determined whether the polymer or core fluid components or bacterial lysates used in the formulation induced gastrointestinal (GI) inflammation or toxicity in mice (Fig. 5A). For this, we orally gavaged six week-old C57BL/6J male and female mice with the individual components used in our droplet formulations or 3% dextran sodium sulfate (DSS) as a positive control, since it is known to induce colitis [30, 31]. We measured body weight changes over time, colon lengths, occult bleeding, and examined intestinal histology after euthanasia at day 6. Both male and female mice that were given 3% DSS exhibited significant body weight loss after six days (Fib. 5B-E) while mice treated with CFE buffer, CFE lysate, or polymer components steadily gained weight. Closer examination of H&E stains of the gut epithelium revealed increased immune cell infiltration and intestinal crypt damage in the 3% DSS treated mouse group which was absent in the other mice (Fig. 5F-G). Notably, this effect was more pronounced in male mice. Similarly, we observed significant shortening of the colons in cohorts of 3% DSS treated mice compared to most formulation components, except for the CFE lysate reagents in female mice where we did not observe a difference (Supp. Fig. 3A-B). Lastly, we detected occult bleeding in mouse fecal samples in 3% DSS treated groups while all other mice were negative (Supp. Fig. 3C). To evaluate overall disease severity, the disease activity index (DAI) based on body weight loss, colon lengths, diarrhea and occult blood was determined. High DAI scores were exclusively observed only in DSS-treated mice (Supp. Fig. 3D), supporting the usage of this formulation in oral delivery applications.

**Figure 5.**
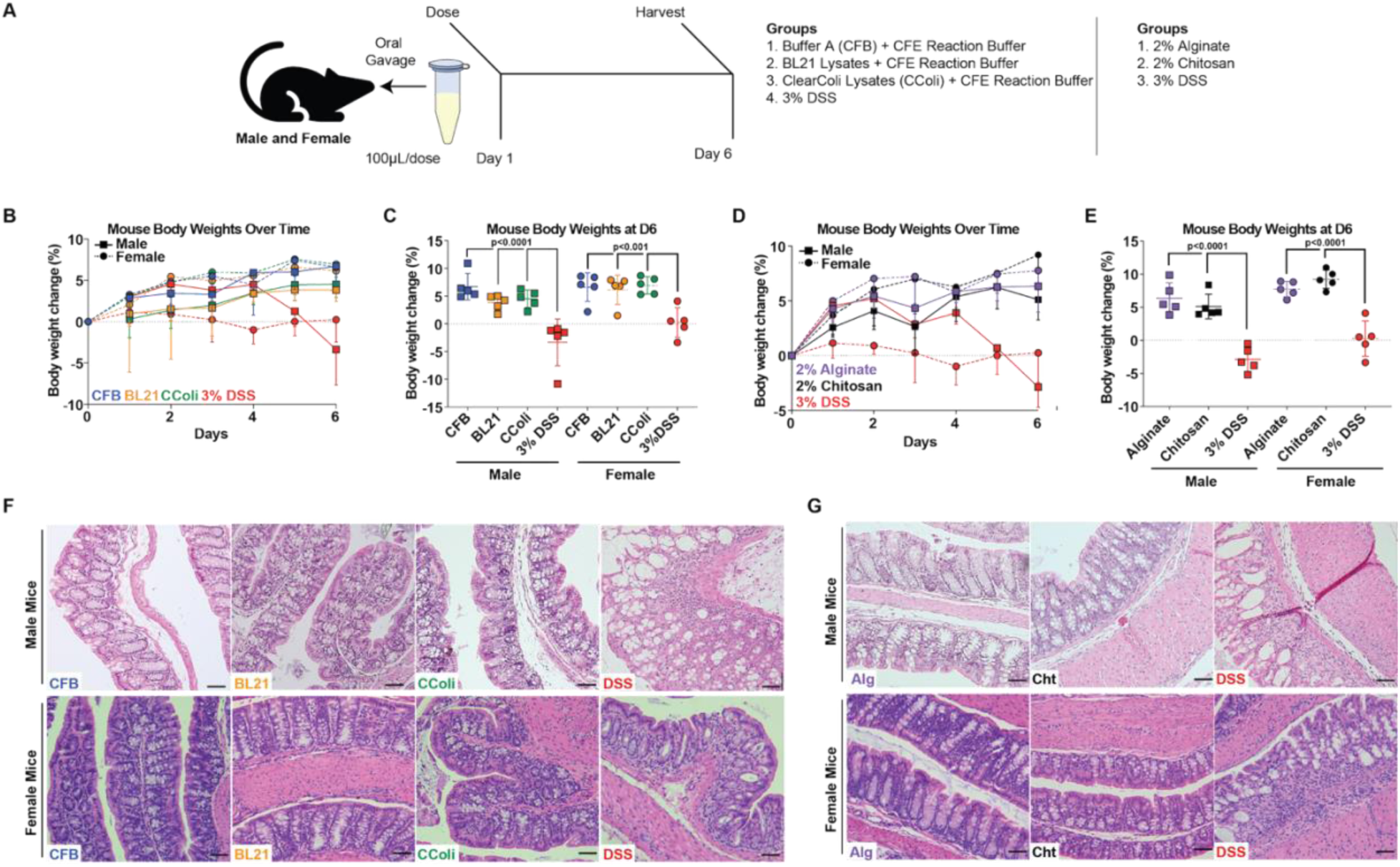
Cell-free lysates and encapsulation components do not induce inflammation in mouse gastrointestinal tracts. (**A**) Schematic depicting dosing of mice with cell-free and encapsulation components. (**B**) Line graph depicting changes in body weights of mice over time after oral administration of cell-free components. n=5 mice per group. (**C**) Dot plot of change in mouse body weights at day 6 after oral administration of CFE components. (**D**) Line graphs depicting changes in body weights of mice over time after oral administration of alginate or chitosan polymers. n=5 mice per group. (**E**) Dot plot depicting changes in body weights of mice at day 6 after oral administration of alginate or chitosan polymers. (**F**) Micrographs of intestinal sections from male (top) and female (bottom) mice after exposure to CFE components stained with H&E (**G**) Micrographs of H&E-stained intestinal sections from male (top) and female (bottom) mice after exposure to alginate or chitosan polymers. Scale bar=100µm.

### Alginate/chitosan microcapsules exhibit controlled delayed release in mouse GI tracts

Since our data suggested that Alg/Cht microcapsules could release their cargo in simulated GI conditions *in vitro* and that individual components were not inflammatory or toxic to mice, we wanted to next track cargo delivery *in vivo*. However, we were concerned that delivery of a fluorescent protein into the mouse GI tract would not be compatible with visualization due to dilution and possible proteolytic degradation after release. To circumvent this issue, we loaded fluorescent cargo consisting of 0.2µm red fluorescent beads, FITC-dextran (70,000 MW), and DAPI into Alg/Cht microcapsules to facilitate tracking before orally gavaging mice and examining their GI release over time (Fig. 6A). We could not detect significant FITC or specific DAPI signal within the tissues, likely due to the large volume dilutions encountered after delivery and cargo release (not shown). However, we could detect the 0.2µm fluorescent beads and collect them by washing out sections of the mouse GI tracts for imaging and flow cytometry. Compared to the unencapsulated cargo alone, where the fluorescent beads were localized to the ceca and colon by 3h post-gavage (Fig. 6B), we observed a delayed transit through the GI tracts of mice orally gavaged with the Alg/Cht encapsulated cargo (Fig. 6C). The cargo was largely absent from the stomach, showing accumulation in the upper and lower small intestine by 3h before passing to the ceca and colon by 6h post-gavage, suggesting a steady controlled passage and release through the mouse GI tract.

**Figure 6.**
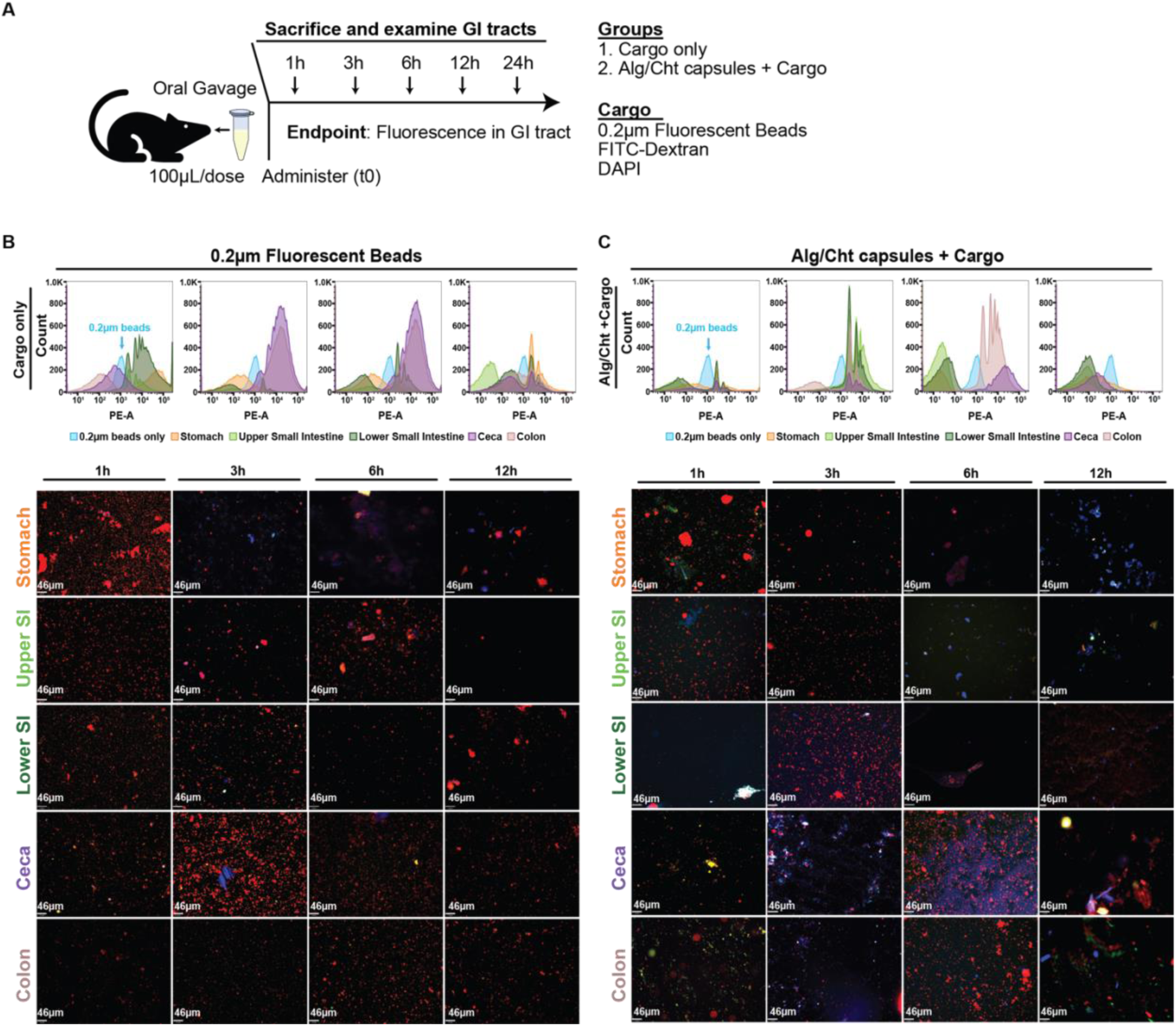
Alg/Cht microcapsules exhibit delayed release of encapsulated cargo in the mouse GI tract. (**A**) Schematic of oral dosing regime for Alg/Cht microcapsules in mice. (**B**) Flow cytometry plots showing 0.2µm fluorescent beads collected from different sections of the mouse GI tract (top), along with representative fluorescence micrographs (bottom). Scale bar=46µm.

## Discussion

Oral delivery of therapeutics is enticing due to the ease of use. However, oral delivery of protein-based therapeutics is challenging as they must be produced, purified and after delivery, must survive the harsh proteolytic conditions of the stomach. Cell-free protein expression enables rapid, decentralized production of proteins, which could be adapted for oral application. However, this remains a challenging task as protein therapeutics are sensitive to their local environment, which may cause their denaturation, aggregation, degradation and loss of activity. This necessitates the use of safe delivery platforms that can insulate and protect protein cargo, such as alginate and chitosan-based systems which remain largely unexplored for these applications.

In this work, we demonstrated that CFE lysates that produce mCherry protein can be encapsulated within Alg/Cht microcapsules and exhibit favorable stability and release characteristics. The fluorescence emitted by mCherry presented a suitable readout of environmental protection, as its chromophore is susceptible to protonation and loss of fluorescence at low pH [32, 33]. Acetic acid was able to penetrate the W/O droplets and inhibit mCherry protein fluorescence during crosslinking, although it recovered after removal of the low pH conditions (Supp. Fig. 1A). In contrast, fully crosslinked Alg/Cht encapsulated mCherry protein retained its activity throughout the *in vitro* release assay, even during incubation in SGF for an hour at 37°C with vigorous mixing (Fig. 4B-C and 4E-F), suggesting that the protein was protected from exposure to the acidic environment. Notably, although we did not include ammonium salts in the SGF formulation [29], reasoning that it would have a minimal effect on droplet stability compared to the low pH, proteases, and other salts in solution, we observed similar behavior of Alg/Cht droplets using mouse GI fluid extracts, demonstrating controlled cargo release *in vitro*.

Crucially, we also observed similar behavior from Alg/Cht droplets *in vivo* after oral gavage, where minimal release of 0.2µm fluorescent beads was observed in the mouse stomach. While the unencapsulated fluorescent beads rapidly passed through the GI tract and settled in the ceca and colon within 3h post-gavage, the encapsulated fluorescent beads exhibited a more controlled and delayed release where fluorescence was strongly detected in the small intestines at 3h before passage to the ceca and colon by 6h (Fig. 5B-C). However, transit time through the GI tract and release kinetics could be variable when comparing protein vs. fluorescent bead cargo as the fluorescent beads are much larger than typical proteins. Still, our *in vitro* release data would suggest that the droplets remained stable at least through the stomach environment and that cargo release mainly occurs in the small intestine, which is ideal for protein-based vaccines since the small intestine harbors organized GALT and antigen sensing and sampling mechanisms for induction of mucosal immune response [34].

Importantly, we did not detect any signs of inflammation in the GI tract or other adverse effects in mice orally gavaged with the individual CFE or polymer components. Considering that for systemic administration, the presence of even minute amounts of bacterial lipopolysaccharides (LPS) in recombinant protein preparations can be a major concern due to the pro-inflammatory and pyrogenic properties of LPS, we tested the safety of LPS^high^ BL21 and LPS^low^ ClearColi-derived *E. coli* CFE lysates delivered per orally. We observed no discernible ill effect in mice in either case. This was expected as oral administration of relatively high doses of LPS have been reported to cause no toxicity or systemic inflammation in mammals, birds, or fish [35–38]. We did observe overall less severe colitis in female mice than male mice, even with 3% DSS. However, this is likely due to the protective anti-inflammatory and tissue-repairing effect of estradiol mediated via estrogen receptor β (Erβ) signaling [39, 40] in female mice, and not due to formulation differences. Significantly, these results suggest that crude protein therapeutics are compatible with oral delivery and could potentially be produced and administered locally at the point of use through CFE, without the need for sophisticated purification procedures and equipment.

While we originally envisioned an all-in-one system where CFE lysates could be loaded into Alg/Cht microcapsules and produce proteins *in situ*, our droplet generation process was evidently incompatible with this idea. The acidification step needed to activate internal gelation through release of Ca^+2^ from particulate CaCO_3_ also seemed to irreversibly inhibit protein synthesis, likely through damage to the transcription and translation components present in the extract [41]. Still, the CFE system did tolerate small amounts of acetic acid (Fig. 1B), and some CaCO_3_ seemed to dissolve at that concentration (Supp. Fig. 1B), suggesting that this approach may still be compatible with internal gelation. Interestingly, nano formulations of CaCO_3_ have been reported that significantly reduce the gelation time of alginate [42] due to its more uniform distribution and accessibility within droplets, presenting a more optimal replacement for the larger CaCO_3_ particles used in this study. The quicker gelation implies that reduced acid concentrations could still achieve similar gelation results, potentially retaining more CFE lysate activity within crosslinked droplets, and remains to be explored in further studies.

Nevertheless, we developed Alg/Cht oral delivery vehicles compatible with purified or crude lysate protein cargo that could protect its contents during stomach passage for a delayed release in the mouse GI tract. These properties are essential for oral therapeutics and could be especially useful for oral vaccine delivery. Even though we saw decreased yield from encapsulated CFE lysates, this does not preclude usage for vaccine production as typically smaller amounts of antigens are required for immunization than for treatment. The current vaccine regimes require centralized production, purification, and administration, and are less compatible with a rapid response to emerging disease outbreaks or application in austere environments. Combining CFE systems with oral delivery could enable decentralized vaccine production and faster responses, only requiring suitable DNA templates for protein antigen synthesis, and help democratize access to these critical healthcare resources.

## Materials and Methods

### Plasmids and Sequences

The mCherry gene containing a hexahistidine tag and thrombin cleavage site at the N-terminus was codon optimized for *E. coli* expression before cloning into the pET-15b vector using the NcoI and BamHI restriction sites (Genscript).

### Preparation of stock solutions

Stock solutions of sodium alginate and low molecular weight (MW) chitosan were prepared before usage in the polymer screen and droplet generation. A 2% (w/v) sodium alginate stock (Millipore-Sigma, W201502) was prepared by dissolving sodium alginate in UltraPure DNase/RNase-Free Distilled Water (Invitrogen, 10977023) while stirring on a hot plate at 90°C until fully dissolved. A 0.5% w/v stock of low MW chitosan (Millipore-Sigma, 448869) was prepared by dissolving it in 150mM glacial acetic acid while stirring on a hot plate until fully dissolved before titration to pH 6 with NaOH.

A stock solution of acidified oil was made by dissolving 3.5mL glacial acetic acid (Supelco, AX0073-75) in 1L of soybean oil (Kirkland Signature, 71011).

A stock solution of 0.85% w/v NaCl stock solution was made by dissolving 8.5g NaCl (Sigma-Aldrich, S9888) in 1L of MilliQ water. NaCl solution used for washing the microcapsules were not pH adjusted. NaCl solution used for final resuspension of microcapsules before coating was adjusted to pH 6.

A 20x stock solution of CaCO_3_ was made by mixing 0.108g of CaCO_3_ (21061-250G-F, Millipore Sigma) in 10mL UltraPure water and probe sonicated for 4 minutes, 15s on and 15s off.

Human simulated gastric fluid (hSGF) and simulated intestinal fluid (hSIF) were prepared according to Minekus et al., 2014 [29] with slight modifications (Table 1). Pepsin was added fresh immediately before each use. Mouse gastric fluid and intestinal fluid extracts were obtained by separately rinsing the stomach and small intestine from mice after euthanasia twice with 500μL pure water, using a 20G plastic needle. Each rinse was collected separately, and the second wash was used for analysis, since the first wash still contained semi-solid material from the stomach and intestinal contents.

### Macrocapsule generation

Core fluid containing 0.75% (v/v) sodium alginate (final), CFE lysate, reaction buffer, and purified mCherry protein was added dropwise through a 27 ½ gauge needle at regular intervals into a 0.5% (w/v) CaCl_2_/0.5% (w/v) chitosan solution while stirring at room temperature (RT). The droplets were allowed to crosslink and coat for ∼60 seconds after addition of the last droplet and removed one-by-one via pipette and stored in 1mL UltraPure water.

For capsules with core fluid containing mCherry plasmid, the crosslinked droplets were incubated in a Thermomixer C (Eppendorf) at 30°C while shaking at 800rpm for ∼16 hrs before imaging on a glass slide for fluorescence at 520nm on the Odyssey M to check for mCherry protein production. mCherry protein yields were calculated based on comparisons to fluorescence from the droplet standard curve through densitometry analysis using ImageJ FIJI v1.54p.

### Generation of microcapsules and in-droplet protein synthesis

Water-in-oil (W/O) emulsions were generated using the LDC-1 academic test unit using precision cut steel membranes with 20µm pores (Micropore Technologies) where the flow rate determines droplet size, and the protocol was adapted from Ji et al. 2019 [10].

Briefly, all core fluid solutions contained 0.75% sodium alginate (final), 25% (v/v) CFE lysate and 25% (v/v) reaction buffer, 1x CaCO_3_ (0.054% w/v), and either purified mCherry protein or 20µg/mL plasmid encoding mCherry. Core fluids were prepared fresh before every run. A 10mL Luer-lock syringe (Fisher Scientific, 14-823-2A) was used to draw up 3mL of soybean oil with 1% tween-80 to form a “plug” followed by the core fluid and kept vertically to keep the core fluid at the bottom of the syringe. Tubing (3.2 mm inner diameter) was attached to the syringe and connected to the LDC-1 input port and the glass chamber was filled with 50mL of soybean oil with 1% tween-80. A syringe pump (World Precision Instruments, model no. AL-300) in vertical position was used to push the core fluid into the LDC-1 at a flow rate of 0.5mL/min while stirring at 8.04V. Once all the fluid was released from the syringe, the stirring was stopped and the oil emulsion was transferred to a 250mL beaker on a stir plate while stirring at 375rpm. Acidified soybean oil containing 3.5% (v/v) glacial acetic acid was added to the W/O emulsion to release Ca^+2^ from CaCO_3_ and crosslink the droplets over 15 min. Emulsions were broken by addition of 15mL 1% tween-80 into the beaker. The beaker was left stirring at a lower speed (200rpm) for 30 minutes before transferring the crosslinked droplets to a 50mL Falcon tube and centrifuged at 1500 x g for 10 min at 4°C.

After centrifugation, the oil and tween supernatant was decanted before washing the droplet pellet twice with 5mL 1% tween-80 and centrifuged at 1500 x g for 10 min at 4°C. The droplets were subsequently washed two times in 0.85% NaCl following the same protocol. The wet mass of the droplets was measured after decanting the second wash. Droplets were resuspended to 2mg/mL in 0.85% NaCl, pH 6 and coated with 0.05% low MW chitosan, pH 6 (final) solution for a final droplet concentration of 0.3mg/mL.

For in-droplet protein synthesis, purified mCherry was replaced with 20µg/mL plasmid encoding mCherry and the droplet generation process was suspended after transferring the initial W/O emulsion to the beaker. The emulsion was left to stir overnight on the stir plate at 30°C and 375rpm before proceeding with crosslinking and the rest of the capsule generation protocol.

In-droplet mCherry production from the CFE reaction and incorporation of purified mCherry into droplets was confirmed through fluorescence microscopy using an EVOS M7000 Imaging System (ThermoFisher) and quantified through densitometry analysis of the Coomassie stained SDS-PAGE gel using ImageJ FIJI v1.54p

### Cell-free lysate production

Clear Coli cell-free extracts were prepared from ClearColi BL21(DE3) DUOs Electrocompetent Cells (BioSearch Technologies, 60810-1). The bacteria were streaked for isolation on a Luria (LB) agar and a single colony was used to inoculate a 50mL starter culture in 2x YT medium (Sigma-Aldrich, Y2377) supplemented with 0.5% NaCl and grown overnight in a baffled flask. The starter culture was used to inoculate 2L of pre-warmed 2x YT with 0.5% NaCl to a final optical density (OD_600_) of 0.1-0.12. T7 polymerase expression was induced using 1mM 0.2µm filtered Isopropyl-β-D-thiogalactoside (IPTG; Roche, 11411446001) at OD_600_=0.5-0.8. The ClearColi cultures were subsequently grown until they reached an OD_600_=2.5-2.7 before harvesting.

ClearColi cultures were split across multiple 450mL Nalgene PPCO centrifuge bottles (Thermo Scientific, 3141-0500PK) and pelleted at 8000rcf for 10min at 4°C in a Sorvall RC-5C Plus Superspeed Centrifuge using a Sorvall SLA-3000 rotor and resuspended in 20mL Buffer A (10mM Tris base, 14mM magnesium glutamate, 60mM potassium glutamate (Sigma-Aldrich, T1503, 49605, and G1501 respectively), and 1mM fresh dithiothreitol (DTT; Roche, 10197777001) per pellet. All pellets were combined after resuspension, centrifuged at 8000rcf for 10min at 4°C, and resuspended to a final concentration of 1g/mL Buffer A before freezing at –80°C overnight.

The frozen bacterial slurry was thawed on ice before lysis using a qSonica Q500 probe sonicator with a ¼ inch probe (qSonica, 4420) and 20s on/30s off sonication interval at an amplitude of 24-25% and a final energy input of 3.5-4.5kJ. 1mM DTT was added to the lysed suspension and 1mL aliquots of the suspension were centrifuged at 18,000 x g for 10 minutes at 4°C in an Eppendorf 5417R tabletop centrifuge. The supernatant was collected and incubated in an Eppendorf Thermomixer C at 37°C while shaking at 800rpm for 30 minutes to perform the run-off reaction. Afterwards, the lysates were centrifuged at 10,000 x g for 10 minutes at 4°C and the final supernatant was collected and stored at –80°C.

### Cell-free gene expression (CFE) reaction setup and polymer screen

CFE reactions producing mCherry were set up using a mixture of 20µg/mL pET15b mCherry plasmid DNA template (4.72nM), 25% (v/v) cell-free lysate, 25% (v/v) PANOx-SP reaction buffer [43] with slight modifications (final concentrations: 1.2 mM ATP; 0.86 mM GTP, UTP, and CTP; 170 μg/ml *E. coli* tRNA; 2 mM of each amino acid (-glutamic acid), 0.33 mM NAD; 0.27 mM Acetyl-CoA; 1.5 mM spermidine; 1 mM putrescine; 175 mM potassium glutamate; 10 mM ammonium glutamate; 2.7 mM potassium oxalate; 10 mM magnesium glutamate; and 33 mM PEP), and UltraPure DNase/RNase-Free Distilled Water (Invitrogen, 10977023) to fill the rest of the reaction volume.

Polymer additives were included in the CFE reactions at increasing concentrations (v/v) in 50µL scale CFE reactions to examine their effects on mCherry production. All reactions were incubated at 30°C while shaking at 800rpm for 16hr in a Thermomixer C (Eppendorf) before fluorescence imaging in an Odyssey M scanner (LICORBio, ODM-0288). Fluorescence scans were analyzed using ImageJ FIJI v1.54p.

Larger scale 1mL CFE reactions were performed following the standard component ratios described above to produce mCherry protein for purification. The 1mL reactions were incubated in 12-well plates (Corning, CLS351143) at 1mL per well in a humidity-controlled chamber in a New Brunswick C24 incubator shaker at 30°C 300rpm for 16-18hr.

### Purification of mCherry

After production in 1mL scale CFE reactions, mCherry protein was purified using gravity column immobilized metal affinity chromatography (IMAC). 1mL of Ni^+2^-NTA cOmplete His Tag Purification Resin slurry (Roche, 5893682001), equivalent to 500µL of packed beads, was loaded onto a 12mL Bio-Spin disposable chromatography column (Bio-Rad, 7326008) and equilibrated with six packed-bead volumes of equilibration buffer (10 mM imidazole, 50 mM NaH2PO4, and 300 mM NaCl, pH 8). All subsequent buffers used in the purification process differed only in imidazole concentrations. The CFE reaction mixture was added to the column and incubated at 4°C while mixing for 1hr. After incubation, the column was washed with 12 packed-bead volumes of wash buffer (20 mM imidazole) before elution with 3.5 packed bead volumes of elution buffer (250 mM imidazole). A final elution of 0.6 packed bead volumes of high imidazole elution buffer (500 mM imidazole) was performed to ensure complete elution of mCherry. Peak mCherry-containing fractions were determined by fluorescence, pooled, and concentrated using Amicon® Ultra Centrifugal Filter, 10 kDa MWCO (Millipore-Sigma, 5010) to >1mg/mL. mCherry protein was then dialyzed (two rounds) into 1x phosphate buffered saline, pH 7.4 (Gibco, 70011069) in 6-8 kDa MWCO dialyzers (Millipore-Sigma, 75104-M) at a PBS volume 1000x the protein volume. Upon completion of dialysis, the purified mCherry protein was removed from the dialysis chamber, quantified via BSA standard curve on an SDS-PAGE gel, and stored at –20°C for future use.

### SDS-PAGE and western blot

SDS-PAGE and western blot analysis was performed for CFE produced proteins inside and outside of microdroplets. Samples were mixed with 4x NuPAGE™ LDS Sample Buffer and 10x NuPAGE™ Sample Reducing Agent (Invitrogen, NP0007 and NP0009 respectively) before denaturing at 95°C for 5 minutes and loaded onto NuPAGE™ 4 to 12% Bis-Tris protein gels (Invitrogen, NP0323BOX) and run at 200V for 35min in 1x NuPAGE MES SDS Running Buffer (Invitrogen, NP0002). 1µL of total CFE reactions from the polymer screen was loaded onto gels. From the *in vitro* release assays, 1µL of droplets or 9.75µL of droplet supernatant were used. Gels were either stained for total protein using Coomassie stain from the GenScript eStain L1C Protein Staining Kit (GenScript, L00753) or used for western blot transfer.

For western blots, proteins were transferred onto PVDF membranes (Invitrogen, IB24001) using the iBlot 2 Gel Transfer device (Invitrogen, IB24001) using the default P0 protocol. The membrane was blocked in 5% w/v blotting grade blocker (Bio-Rad, 1706404XTU) in 1x PBS-T (1x PBS pH 7.4, 0.1% Tween 20) for 1 hour at RT while rocking and then incubated in primary antibody (1:1000 for mouse anti-(H)_5_ antibody (Qiagen, 34660) or 1:2000 mCherry Rabbit monoclonal antibody (Cell Signaling Technology, 43590S) in 5% blocking buffer) overnight at 4°C. Primary antibody was removed by washing the membrane in 1x PBS-T four times for 5 minutes each at RT, incubated with secondary antibody in blocking buffer (1:10,000 for VRDye 490 Donkey Anti-Rabbit IgG secondary antibody (LICORbio, 926-49013) and IRDye 800CW Donkey Anti-Mouse IgG secondary antibody (LICORbio, 926-32212)) for 1 hr at RT, and washed again four times for 5 minutes each in 1x PBS-T before imaging on the LI-COR Odyssey M imager (ODM-0288).

### In-vitro cargo release testing

To test cargo release *in vitro*, 250µL of microdroplets were centrifuged at 1500 rcf for 5 min, decanted, and resuspended to 0.3mg/mL in hSGF. The droplets were incubated in a Thermomixer C at 37°C while shaking at 800rpm for 1 hr before a 50µL aliquot was removed for downstream analysis. hSIF was added to the remaining droplets in a 1:1 (v/v) hSGF:hSIF ratio. The droplets were further incubated under the same conditions and timepoint aliquots were collected at 30 min, 1 hr, 4 hrs, and 20 hrs after addition of hSIF.

After collection, 1µL of all aliquots were separated for fluorescence microscopy before the rest of the sample was centrifuged at 1500 rcf for 5 min at 4°C. The supernatant was removed into a different 1.5 mL tube after centrifugation and both supernatant and droplet pellets were placed in ice to slow release. After the end of the time series experiment, all tubes were imaged on the Odyssey M and densitometry analysis performed using ImageJ FIJI v1.54p. Release testing using mGF and mIF followed identical procedures.

### In-vivo release testing

To test cargo release from Alg/Cht droplets in the GI tract, we used 0.2% v/v red fluorescent beads (580/605, 0.2µm) (Invitrogen, F8763), 10mg/mL DAPI and 1mg/mL FITC-dextran as cargo. Mice were fasted for 6hrs prior to PO administration of droplets containing cargo or unencapsulated cargo only. Mice were euthanized at 1, 3, 6, and 12 hours after oral gavage and the GI tract was excised and stomach, upper SI, lower SI, cecum and colon were separated. Each GI section was washed with 1mL PBS with 0.02% NaN3. Ceca and colon washes were centrifuged at 3000 rpm for 10 minutes to remove fecal material. Samples were then analyzed by flow cytometry using the BD FACSSymphony A1 and examined by fluorescence microscopy with the Leica DM4000 B (Leica Microsystems) equipped with a QIClick digital CCD camera (QImaging). Flow cytometry data were processed using FlowJo v10.10.0, and acquired images were analyzed using Volocity Software 7.0.

### Animals and housing

For *in vivo* studies, 6– to 8-week old male and female C57BL/6J mice were used and obtained from The Jackson Laboratory (Bar Harbor, ME).

### Assessment of the safety of CFE components and polymers

The safety of CFE components and encapsulation polymers namely, Buffer A, BL21 lysate, ClearColi lysate, Alginate 2%, and Chitosan (2%) were examined *in vivo*. All CFE reagents contained PANOx-SP reaction buffer. For this, mice were fasted for 6h before administration of 100 mL of test solution via gastric gavage using 20G x 38mm (0.603mm inner diameter) curved gavage needles. As a positive control for intestinal inflammation and weight loss, 3% dextran sodium sulfate (DSS, MP Biomedicals, Solon, OH, USA) was administered in the drinking water. The DSS solution was freshly prepared daily to prevent precipitation. Following administration of the test formulations or DSS, mice were monitored daily for changes in body weight, overall appearance, behavior (piloerection, hunched posture, mobility, and excessive huddling), stool consistency, and the presence of blood in stool or in rectal swabs. On day 6, mice were euthanized and the GI tracts were excised to measure colon lengths. The colons and small intestines (SI) were then gently rinsed to remove chyme and fecal pellets before processing for histological analysis. Fecal occult blood was assessed using a commercial kit (Beckman Coulter, Brea, CA, USA), and the disease activity index was calculated based on body weight loss, diarrhea, and occult blood or gross bleeding [44].

### Histological examination

To examine their entirety, colon and small intestines were prepared as “Swiss-rolls” and fixed overnight in 4% formaldehyde for histology. Specimens were then embedded in paraffin and thin tissue sections were then stained with H&E stain, followed by a histological analysis focusing on inflammatory cell infiltration and changes in intestinal tissue architecture.

### Ethics Statement

All procedures involving animals received prior approval from the Southern Illinois University Institutional Animal Care and Use Committee (IACUC) protocol #24-023 and the Animal Care and Use Review Office (ACURO, protocol CB11447.e001) within the USAMRDC Office of Research Protections (ORP). The study was carried out in strict compliance with the National Institutes of Health (NIH) guidelines, the Animal Welfare Act, and all relevant U.S. federal regulations.

## Author Contributions

Investigation: F.M., F.L., D.S., E.W., K. L., V.K., and S.H.-P. Methodology: F.M., F.L., V.K., and S.H.-P. Formal Analysis: F.M., F.L., D.S., V.K., and S.H.-P. Conceptualization: V.K., M.A.C., and S.H.-P. Funding Acquisition: V.K., M.A.C., and S.H.-P. Supervision: V.K., and S.H.-P. Writing – original draft: F.M., F.L., V.K., M.A.C., and S.H.-P. Writing – review and editing: F.M., F.L., V.K., M.A.C., and S.H.-P.

## Funding

This work was supported by the Defense Threat Reduction Agency [project number HDTRA1447641].

## Conflicts of interest

The authors declare no conflicts of interest from this work.

## Supporting information

Table 1. Human simulated gastric and intestinal fluid formulations

## Acknowledgements

We would like to thank M. Lyman for the mCherry plasmid and C. Liu, E. Laurence, C.W. Ye, and N. N. Watkins for their helpful discussions. Work at the Lawrence Livermore National Laboratory was performed under the auspices of the U.S. Department of Energy under Contract DE-AC52-07NA27344.

---

**Figure S1.**
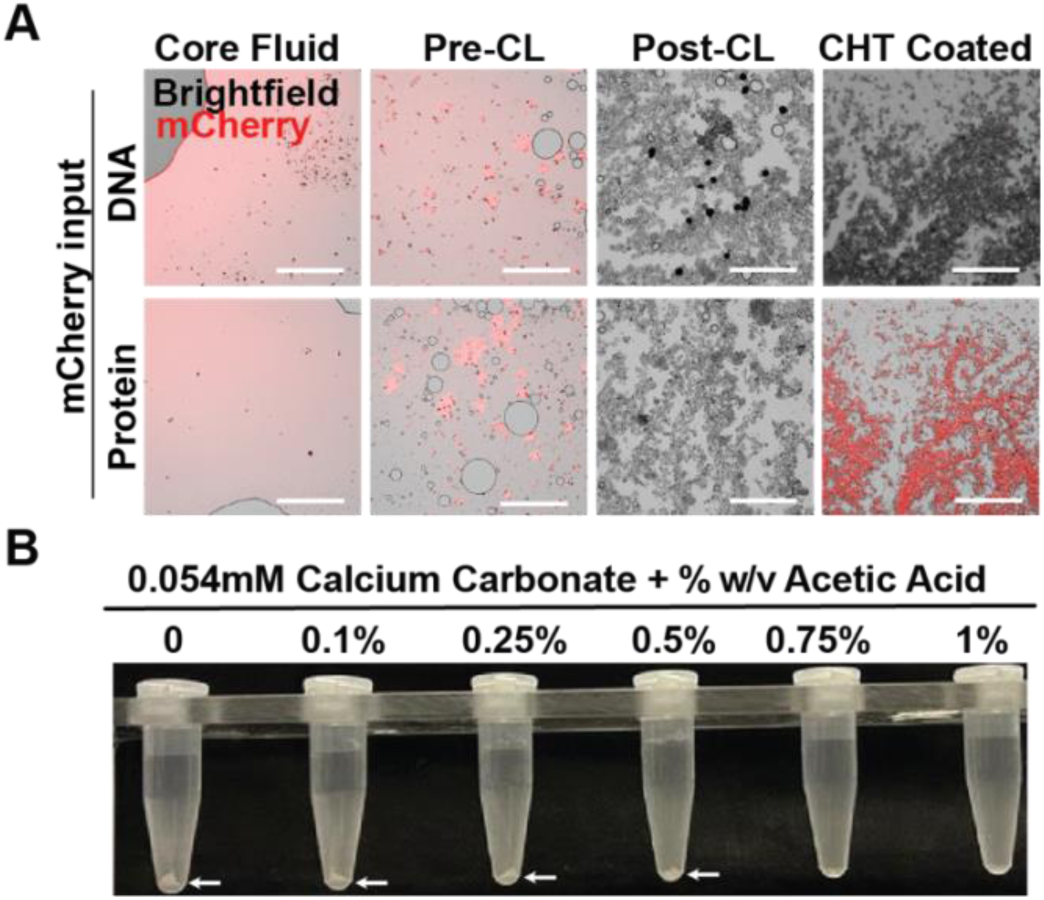
Acetic acid concentrations sufficient to release Ca^+2^ in internal gelation inhibit cell-free gene expression, related to Figure 3. (**A**) Fluorescence micrographs of CFE mCherry (top) and purified mCherry (bottom) sampled from different steps during droplet generation. Scale bar=275µm. (**B**) Photograph of tubes containing 0.054mM CaCO_3_ solubilized with increasing concentrations of acetic acid.

**Figure S2.**
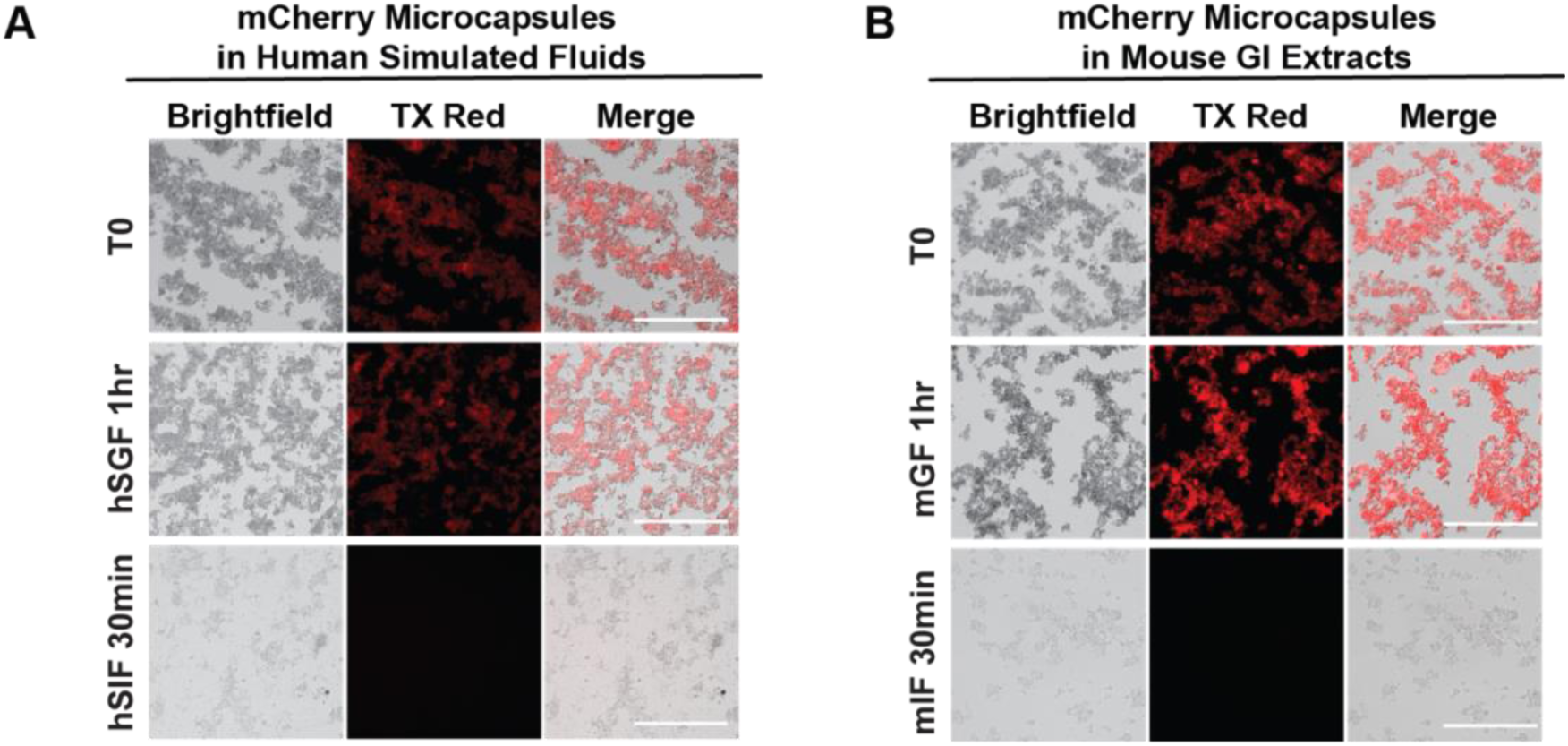
mCherry protein from Alg/Cht microcapsules is released after exposure to intestinal-like conditions. (**A**) Fluorescence micrographs of mCherry-loaded Alg/Cht microcapsules during *in vitro* release assay using human simulated gastric and intestinal fluids. (**B**). Fluorescence micrographs of mCherry-loaded Alg/Cht microcapsules during *in vitro* release assay using mouse gastric and intestinal extracts. All scale bars=275µm.

**Figure S3.**
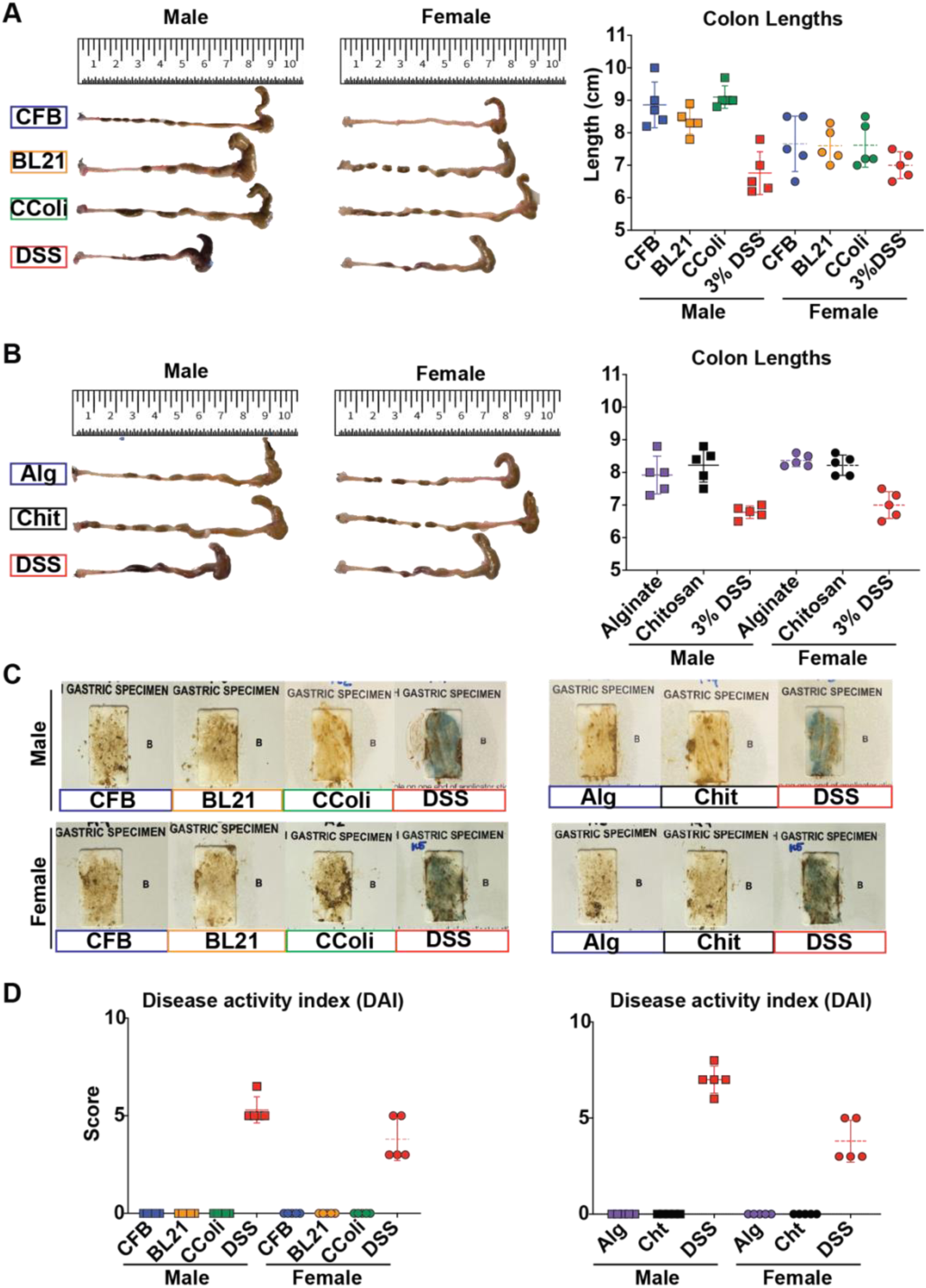
Male and female mouse gastrointestinal tracts show no adverse effects from encapsulation materials, related to Figure 5. (**A-B**) Photographs of colons (left) collected from male or female mice and dot plots depicting colon lengths (right) of mice orally gavaged with cell-free (A) or polymer components (B). (**C**) Representative photographs of occult bleeding assay from male and female mice. A positive result (blue color on cards) indicates blood presence. (**D**) Dot plot showing calculated disease activity index scores.

## Notes

### Competing Interest Statement

The authors have declared no competing interest.

